# Comparison of gnotobiotic communities reveals unexpected amino acid metabolism by the pre-weaning microbiome

**DOI:** 10.1101/2024.02.22.581588

**Authors:** Jean-Bernard Lubin, Michael A. Silverman, Paul J. Planet

**Author notes:** Co-senior authors (Michael A. Silverman, Paul J. Planet).

## Abstract

The intestinal microbiome during infancy and childhood has distinct compositions and metabolic functions to that of adults. We recently published a gnotobiotic mouse model of the pre-weaning microbiome (PedsCom), which retains a pre-weaning configuration during the transition from a milk-based diet to solid foods and leads to a stunted immune system and susceptibility to enteric infection. Here we compared the phylogenetic and metabolic relationship of the PedsCom consortium to the adult-derived gnotobiotic communities, Altered Schaedler Flora and Oligo-MM12. We find that PedsCom contains several unique functions relative to adult-derived mouse consortia. In particular, amino acid degradation metabolic modules are more prevalent among PedsCom isolates, which is in line with the ready availability of these nutrients in milk. Indeed, metabolomic analysis showed significantly lower levels of free amino acids in the intestinal contents of adult PedsCom colonized mice versus Oligo-MM12 controls. Thus, enhanced amino acid metabolism is a prominent feature of the pre-weaning microbiome that may facilitate design of early life microbiome interventions.

## Introduction

The most dramatic physiologic changes in the mammalian microbiome occur at weaning^1^. The transition from milk to a solid food-based diet is accompanied by radical shifts in intestinal microbiome diversity and composition, as post-weaning associated microbes take advantage of newly available nutrient sources. Disruptions to the early-life microbiome that occur prior to and during weaning have detrimental impacts on host immune system development, increasing the risk of allergy and autoimmunity later in life^2,3^. To better understand the physiologic impacts of the early-life microbiome, we developed a nine-member consortium (PedsCom) to model the unique composition and function of mammalian pre-weaning intestinal microbiomes^4^. We found that restricting gnotobiotic mice to a pre-weaning community in adulthood stunted key markers of immune system maturation (IgA and peripheral regulatory T cells), indicating that development of the microbiome during weaning is crucial for normal immune system maturation. Additionally, the PedsCom consortium was resistant to diet-induced shifts in community structure during weaning. This intriguing stunting of the microbiome raises multiple questions about the metabolic capabilities of PedsCom, including which nutrients drive the critical shifts in microbial composition.

The change in carbohydrates during the transition from milk to a solid food-based is often invoked as the key driver of weaning associated changes in the intestinal microbiome. The carbohydrate fraction of milk is primarily composed of lactose and host-indigestible milk oligosaccharides, while solid foods typically contain primarily starches and host-indigestible fiber. Although lactose is a major source of available energy in the pre-weaning gut, the majority of this nutrient is consumed and utilized by the host in the small intestine^5^. As such, alternate milk-derived carbohydrates such as human milk oligosaccharides (HMOs) likely have more impact on the pre-weaning microbiome in the distal gut. Indeed, HMOs promote neonatal colonization of beneficial species such as *Bifidobacterium* spp.^6,7^.

The microbiome changes associated with lipid and protein nutrient composition between pre- and post-weaning diets are less well understood. Lipids are the second most abundant macronutrient after lactose in human milk and provide ∼40% of the total energy content^8^. The protein fraction of milk is another distinguishing factor of pre- and post-nutrient availability. Nitrogen is considered a limiting nutrient in the colonic microbiome^9^, and there is growing interest in the role of nitrogen sources as driver of microbiome dynamics^10^. Milk proteins consist of different amino acid proportions from non-milk sources, containing much higher proline and glutamine content^11^. Additionally, free amino acids in milk primarily provide glutamic acid, taurine, alanine and glutamine to pre-weaning gut environment^12,13^. The differences in the abundance and composition of all three macronutrients (carbohydrate, lipid, and protein) in milk versus solid food likely contribute to the colonization and metabolic output of pre-weaning microbiota.

Here we compare the phylogenetic diversity and metabolic capabilities of the preweaning gnotobiotic community PedsCom to two gnotobiotic communities [Oligo-MM12 and Altered Schaedler Flora (ASF)] that model the adult microbiome to better understand the microbial and metabolic dynamics that occur during weaning. Using functional predictions from whole genomes, we measured the diversity of metabolic potential, focusing on functions overrepresented in the pre-weaning human microbiome and the nutrients found in human milk. We then interrogated these predictions by comparing the intestinal metabolomic profiles of adult PedsCom- and Oligo-MM12-colonized mice.

## Results

### Phylogenetic distribution of PedsCom isolates reflects the composition of pre-weaning microbiomes

Phylogenetic and taxonomic diversity can be used to predict the metabolic functions of the microbiome. We first compared PedsCom to two established gnotobiotic consortia, ASF and Oligo-MM12, using a phylogenetic approach based on the highly conserved RNA polymerase subunit beta amino acid sequence (RpoB)^14^ (**Fig. 1 and Table 1**). PedsCom contains representatives from three phyla: *Bacillota, Bacteroidota* and *Pseudomonadota*, and ASF includes microbes from *Bacillota, Bacteroidota* and *Deferribacterota*, while Oligo-MM12 is more diverse at the phylum level with the inclusion of the same three phyla in PedsCom as well as the *Actinomycetota* and *Verrucomicrobiota*.

**Table 1.** Bacterial isolates of PedsCom, Altered Schaedler Flora and Oligo-MM12 consortia. Listing of consortium isolates investigated in this study. Taxonomic classification to the family level, genome size, GC content and number of annotated Kegg Orthologs (KOs) included for comparison.

| Consortium | Genome | Phylum | Class | Order | Family | Genome size (MB) | GC content | KO count |
| --- | --- | --- | --- | --- | --- | --- | --- | --- |
| PedsCom | Mammalicoccus sciuri PC04 | Bacillota | Bacilli | Bacillales | Staphylococcaceae | 2.84 | 32.6 | 1644 |
|  | Staphylococcus xylosus PC20 | Bacillota | Bacilli | Bacillales | Staphylococcaceae | 2.83 | 32.8 | 1563 |
|  | Enterococcus faecalis PC15 | Bacillota | Bacilli | Lactobacillales | Enterococcaceae | 2.88 | 37.5 | 1466 |
|  | Lactobacillus johnsonii PC38 | Bacillota | Bacilli | Lactobacillales | Lactobacillaceae | 1.92 | 34.6 | 984 |
|  | Ligilactobacillus murinus PC39 | Bacillota | Bacilli | Lactobacillales | Lactobacillaceae | 2.48 | 39.7 | 1197 |
|  | Clostridium intestinale PC17 | Bacillota | Clostridia | Eubacteriales | Clostridiaceae | 4.69 | 30.4 | 2108 |
|  | Anaerostipes sp. PC18 | Bacillota | Clostridia | Eubacteriales | Lachnospiraceae | 3.39 | 43.5 | 1605 |
|  | Parabacteroides distasonis PC19 | Bacteroidota | Bacteroidia | Bacteroidales | Tannerellaceae | 5.33 | 45.1 | 1535 |
|  | Kosakonia cowanii PC08 | Pseudomonadota | Gammaproteobacteria | Enterobacterales | Enterobacteriaceae | 4.99 | 55.8 | 2886 |
| Altered Schaedler Flora | Lactobacillus intestinalis ASF360 | Bacillota | Bacilli | Lactobacillales | Lactobacillaceae | 1.87 | 35.9 | 861 |
|  | Ligilactobacillus murinus ASF361 | Bacillota | Bacilli | Lactobacillales | Lactobacillaceae | 2.11 | 40.0 | 1125 |
|  | Clostridium sp. ASF356 | Bacillota | Clostridia | Eubacteriales | Clostridiaceae | 2.81 | 30.9 | 1218 |
|  | Eubacterium plexicaudatum ASF492 | Bacillota | Clostridia | Eubacteriales | Eubacteriaceae | 6.10 | 42.9 | 1733 |
|  | Schaedlerella arabinosiphila ASF502 | Bacillota | Clostridia | Eubacteriales | Lachnospiraceae | 6.24 | 47.9 | 1968 |
|  | Pseudoflavonifactor sp. ASF500 | Bacillota | Clostridia | Oscillospirales | Oscillospiraceae | 3.70 | 58.8 | 1274 |
|  | Parabacteroides goldsteinii ASF519 | Bacteroidota | Bacteroidia | Bacteroidales | Tannerellaceae | 6.86 | 43.4 | 1594 |
|  | Mucispirillum schaedleri ASF457 | Deferribacterota | Deferribacteres | Deferribacterales | Mucispirillaceae | 2.34 | 31.2 | 1274 |
| Oligo-MM12 | Bifidobacterium animalis YL2 | Actinomycetota | Actinomycetes | Bifidobacteriales | Bifidobacteriaceae | 2.02 | 60.2 | 919 |
|  | Enterococcus faecalis KB1 | Bacillota | Bacilli | Lactobacillales | Enterococcaceae | 3.03 | 37.2 | 1488 |
|  | Limosilactobacillus reuteri I49 | Bacillota | Bacilli | Lactobacillales | Lactobacillaceae | 2.01 | 38.8 | 1060 |
|  | Blautia coccoides YL58 | Bacillota | Clostridia | Eubacteriales | Lachnospiraceae | 5.13 | 45.7 | 1954 |
|  | Enterocloster clostridioformis YL32 | Bacillota | Clostridia | Eubacteriales | Lachnospiraceae | 6.97 | 48.1 | 2252 |
|  | Acutalibacter muris KB18 | Bacillota | Clostridia | Eubacteriales | Oscillospiraceae | 3.80 | 54.6 | 1296 |
|  | Flavonifactor plautii YL31 | Bacillota | Clostridia | Eubacteriales | Oscillospiraceae | 3.81 | 60.9 | 1558 |
|  | Clostridium innocuum I46 | Bacillota | Erysipelotrichia | Erysipelotrichales | Erysipelotrichaceae | 4.47 | 43.1 | 1621 |
|  | Bacteroides caecimuris I48 | Bacteroidota | Bacteroidia | Bacteroidales | Bacteroidaceae | 4.75 | 42.6 | 1326 |
|  | Muribaculum intestinale YL27 | Bacteroidota | Bacteroidia | Bacteroidales | Muribaculaceae | 3.31 | 50.1 | 1099 |
|  | Turicimonas muris YL45 | Pseudomonadota | Betaproteobacteria | Burkholderiales | Sutterellaceae | 2.89 | 44.1 | 1298 |
|  | Akkermansia muciniphila YL44 | Verrucomicrobiota | Verrucomicrobiae | Verrucomicrobiales | Akkermansiaceae | 2.74 | 55.7 | 1082 |

**Figure 1.**
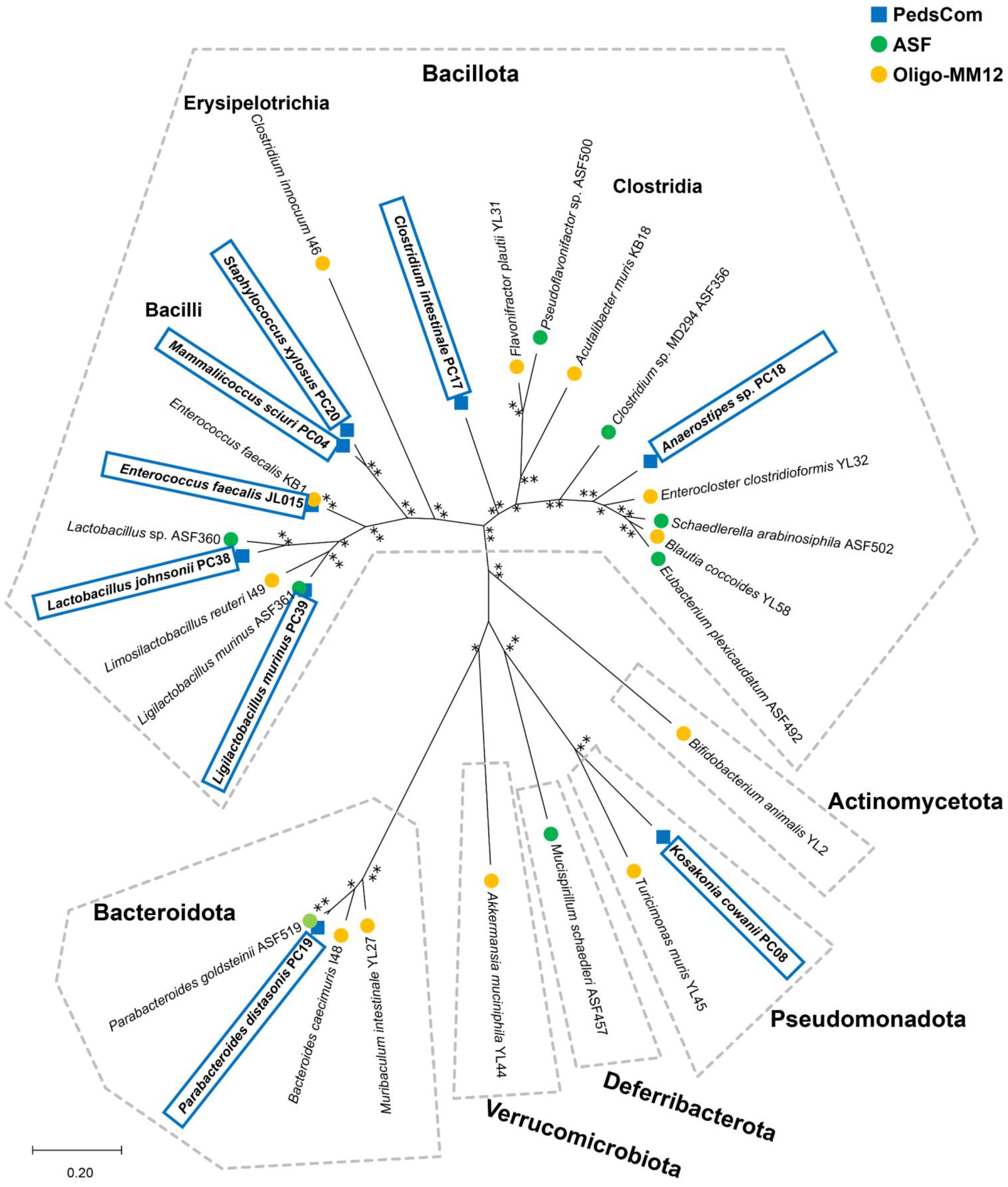
PedsCom consortium models phylogenetic composition of pre-weaning microbiomes. DNA polymerase subunit beta (RpoB) amino acid sequence maximum-likelihood tree of PedsCom, Altered Schaedler Flora (ASF) and Oligo-MM12 isolates. The tree with the highest log likelihood is shown. Percentage of bootstrap trees in which associated taxa were clustered represented by ** > 75%, * > 60%.

The unique presence of the *Pseudomonadota* (represented by *Kosakonia cowanii*, formerly *Enterobacter cowanii*) in PedsCom is reflective of mammalian early-life microbiomes, which contain very high levels of the *Pseudomonadota* family *Enterobacteriaceae. Enterobacteriaceae* are among the first colonizers of the human infant gut^15–17^, and they are major contributors to the pre-weaning microbiomes of mice and humans^18,19^. It has been proposed that facultative anaerobes affiliated with the *Enterobacteriaceae* contribute to the reducing environment needed for colonization of other anaerobes with expanded metabolic capabilities^20,21^. Other metabolic studies have implicated *Enterobacteriaceae* (*E*.*coli*) in increased amino acid consumption in the early-life gut^22^.

The *Bacillota* representatives from the three gnotobiotic communities are distributed into three clades representing the classes *Clostridia, Bacilli* and *Erysipelotrichia*. The *Clostridia* clade primarily includes species of the families *Lachnospiraceae* and *Oscillospiraceae* (formerly known as *Ruminococcaceae*). These families comprise a substantial fraction of the adult gut microbiome and support gut homeostasis^23^. PedsCom contains a single member of *Lachnospiraceae* (*Anaerostipes* sp. PC18), which is in a separate lineage from the representatives in ASF (*Schaedlerella arabinosiphila* ASF502, *Eubacterium plexicaudatum* ASF492) and Oligo-MM12 (*Enterocloster clostridioformis* YL32, *Blautia coccoides* YL58). PedsCom does not contain any members of *Oscillospiraceae* (*Flavonifractor, Pseudoflavonifractor, Acutalibacter spp*.). Instead, PedsCom contains *Clostridium intestinale*, a *Clostridiaceae*, which has been found in higher abundances in human infants relative to adults^24^. *Lachnospiraceae* and *Oscillospiraceae* are well characterized fermenters of host-indigestible fibers into short-chain fatty acids^25^ and are enriched in the intestinal mucosa of the adult murine colon^26^. Taken together, this phylogenetic comparison suggests that critical functions in the adult gut are not recapitulated in PedsCom mice. Indeed, PedsCom mice have much lower levels of short-chain fatty acids in their intestines compared to Oligo-MM12 mice^4^. Thus, the reduced representation of fiber-utilizing bacteria may contribute to PedsCom’s resistance to diet-induced shifts in relative abundance during weaning.

*Bacilli*, such as lactobacilli, staphylococci, and enterococci, are a highly prevalent class of bacteria in the pre-weaning microbiomes of mice and humans^18,19,27^. PedsCom contains two species of *Lactobacillaceae, Lactobacillus johnsonii* and *Ligilactobacillus murinus*. The family *Lactobacillaceae* is highly abundant in the early-life microbiome of mice^28^, and our recent study^4^ found that *L. johnsonii* is more abundant in the small intestine while *L. murinus* dominates the cecum and colon, suggesting niche specialization. *Enterococcus* spp. are also more abundant in gut microbiomes of pre-weaning mice^29,30^, and the PedsCom and Oligo-MM12 *Enterococcus faecalis* isolates are closely related, with an average nucleotide identity (ANI) of 99.1%. PedsCom uniquely contains *Staphylococcaceae* (*Mammaliicoccus sciuri, Staphylococcus xylosus*) which are also more abundant in the pre-weaning microbiomes of mice and humans^19,30^. Overall, the high proportion and relative abundance of *Bacilli* in PedsCom (5 of 9 taxa, average relative abundance of 71% in small intestine, 26% in large intestine at two weeks of age) likely provides many bacterial functions prevalent in the pre-weaning period.

The *Bacteroidota* phylum is prominent in the distal intestines of adult and early-life microbiomes. PedsCom contains *Parabacteroides distasonis*, which is distantly related to the ASF representative *Parabacteroides goldsteinii*, with an ANI of 74.2% that is at the threshold of separate genera^31^. The genome of *P. distasonis* is also 30% smaller than *P. goldsteinii’s* (∼1.6 MB difference), indicating that *P. goldsteinii* likely encodes many functions absent in *P. distasonis*. Oligo-MM12 contains two *Bacteroidota, Bacteroides caecimuris* (*Bacteroidaceae*) and *Muribaculum intestinale* of the family *Muribaculaceae*, also known as S24-7. The Oligo-MM12 *Bacteroidota* are dominant members of the adult mouse intestinal microbiome^32^. In contrast, the relative abundance of *P. distasonis* decreased with age in mice with a complex mature microbial community, analogous with conventionally housed mice^4^. Limiting PedsCom to a single pre-weaning associated *Bacteroidota* likely decreases the fiber-degrading functions found in adult microbial communities. The combination of less fiber-degrading *Lachnospiraceae* and *Oscillospiraceae* and less *Bacteroidota* species in PedsCom likely contributes to lower intestinal short-chain fatty acids levels in adult PedsCom mice.

Overall, the phylogenetic distribution of PedsCom mirrors the pre-weaning microbiome with a high proportion of *Bacilli*, low numbers of fiber-degrading *Clostridia*, and unique representation from the *Enterobacteriaceae*. Together, these taxonomic compositions predict that PedsCom has less ability to metabolize fiber, which is born out in our previous results showing reduced levels of short chain fatty acids^4^.

### Functional potential of PedsCom consortium indicates adaptation to pre-weaning host

To interrogate the phylogenetically inferred metabolic distinctions between PedsCom and the two other communities, we compared the genomes for predicted functional capabilities using the Kyoto Encyclopedia of Genes and Genomes (KEGG) Orthologs (KOs). Briefly, open reading frames in each genome were assigned to a KO using the KEGG automatic annotation server (KAAS) and combined to represent the total predicted metabolic function of each community. A comparison of the specific KOs reveals that PedsCom (4,062) contains a considerably higher number of non-redundant KOs than ASF (2,853) or Oligo-MM12 (3,403) (**Fig. 2A**,**B**).

**Figure 2.**
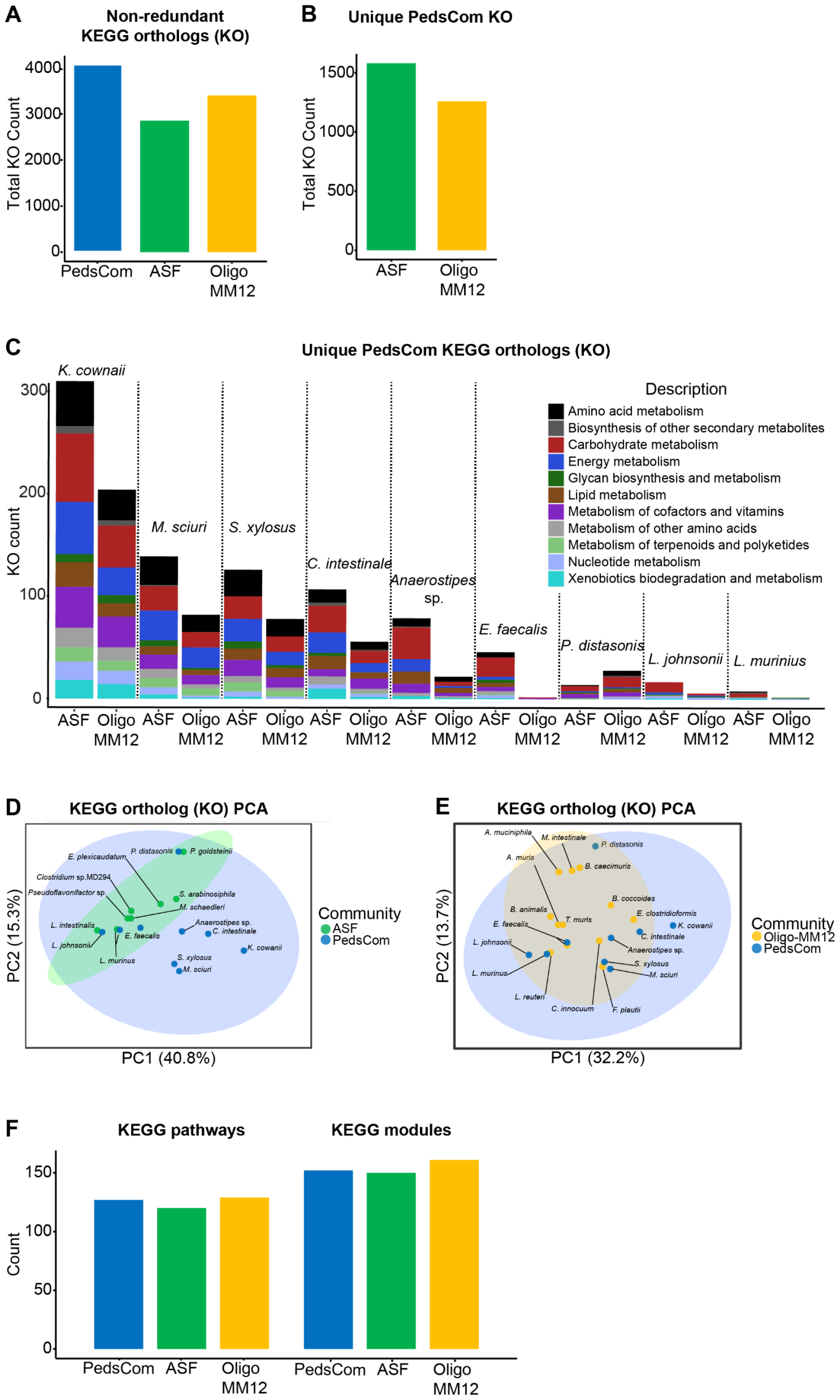
PedsCom consortium isolates members are functionally diverse. **A**.Non-redundant KEGG orthologs (KOs) present in PedsCom, ASF and Oligo-MM12 isolates. **B**.Number of unique KOs present in PedsCom isolates relative to ASF and Oligo-MM12. **C**. Breakdown of unique KEGG orthologs by PedsCom isolate. KEGG orthologs annotated by metabolic function. **D**. Principal component analysis (PCA) plots of KEGG Ortholog pathways of PedsCom and ASF isolates. **E**. Principal component analysis (PCA) plots of KEGG Ortholog pathways of PedsCom and Oligo-MM12 isolates. **F**. Number of non-redundant KEGG pathways and modules present in PedsCom, ASF, Oligo-MM12 consortia.

To determine if the unique KOs present in PedsCom are associated with the functions of the early-life microbiota, we compared the predicted functions of KOs shared by at least three PedsCom isolates with functions overrepresented in human infant microbiomes^17^. Using this approach, we identified several metabolic functions that are (1) overrepresented, and (2) unique to the PedsCom consortium compared to ASF and Oligo-MM12 (**Table 2**). These KOs are associated with glycolysis, gluconeogenesis, the TCA cycle, and metabolism of α-linoleic acid, lipoic acid, galactose, glutathione, ubiquinone/menaquinone, fructose and mannose. Of the PedsCom microbes, *Kosakonia cowanii* is the principal contributor of unique metabolic capability in PedsCom, with smaller contributions from the staphylococcal and clostridial species (**Fig. 2C**). A more global view using principal component analysis (PCA) of KOs assigned to KEGG pathways revealed that PedsCom isolates are predicted to be much less functionally related to each other than ASF isolates (**Fig. 2D**), denoted by the smaller 95% confidence interval that encompasses ASF isolates relative to PedsCom. In agreement with Fig. 2D, these differences in PedsCom are primarily found in the staphylococci (*M. sciuri, S. xylosus), Clostridia* (*Anaerostipes* and *C. intestinale)*, and *K. cowanii* (**Fig. 2C**). The functional diversity of Oligo-MM12 overlapped more with PedsCom, with only *K. cowanii* falling outside the 95% confidence interval of Oligo-MM12 (**Fig. 2E**).

**Table 2.**
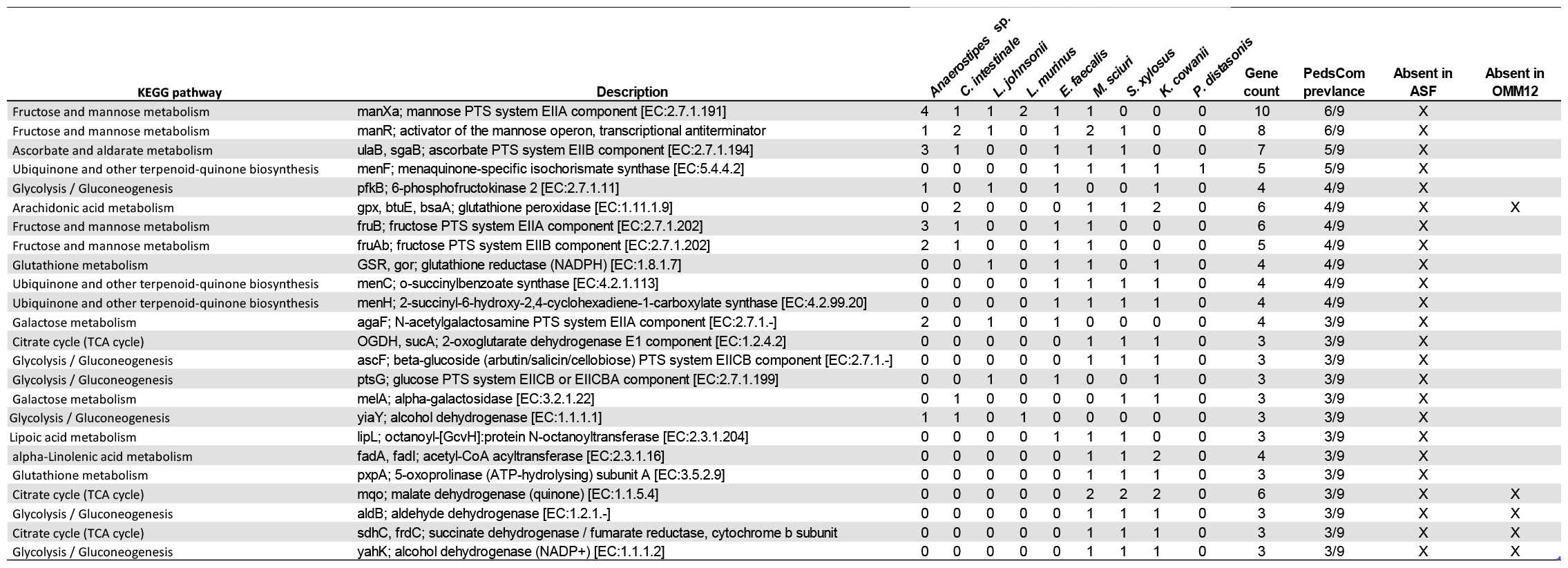
PedsCom unique KOs with functions overrepresented in human infant fecal metagenomes. Most prevalent (shared by ≥ three isolates) Kegg Orthologs (KOs) with functions overrepresented in human infant fecal microbiomes^18^. Only KOs unique to the PedsCom relative to Altered Schaedler Flora and/or Oligo-MM12 are presented.

Despite the increased numbers of KOs in PedsCom, the numbers of KEGG pathways [PC (127), ASF (120), OMM12 (129)] and modules [PC (152), ASF (150), OMM12 (161)] in each consortium were comparable (**Fig. 2F**). These findings of a higher number of KOs in PedsCom, but a similar number of KEGG pathways and modules indicates that the unique KOs present in PedsCom likely perform similar or overlapping metabolic functions. This metabolic ‘redundancy’ in PedsCom genomes may be reflective of uniformly subsisting on milk diet compared to a diverse solid food diet.

### Gut specific metabolic modules reveal a bias towards amino acid degradation in PedsCom isolates

To investigate the gut-specific metabolic potential of PedsCom compared to ASF and Oligo-MM12, we applied a manually curated set of KEGG modules specific for the gut ecosystem called Gut Metabolic Modules (GMMs)^33^. GMMs accurately assign KOs to processes in the gut and avoid inclusion of superfluous functions that are unlikely to contribute to metabolic functions and fitness in the intestine. To determine the presence or absence of each of the 103 GMMs from each consortium we set a threshold of possessing at least 75% of the genes assigned to a GMM. Using this criterion, PedsCom has 69 GMMs while ASF has 51 GMMs and OligoMM12 has 63 (**Fig. 3A**). GMM analysis of PedsCom metabolic potential points to *K. cowanii* as the primary contributor of unique metabolic functions. Of the 19 PedsCom GMMs not found in ASF, 18 of them were found in *K. cowanii*, with nine exclusive to this microbe.

**Figure 3.**
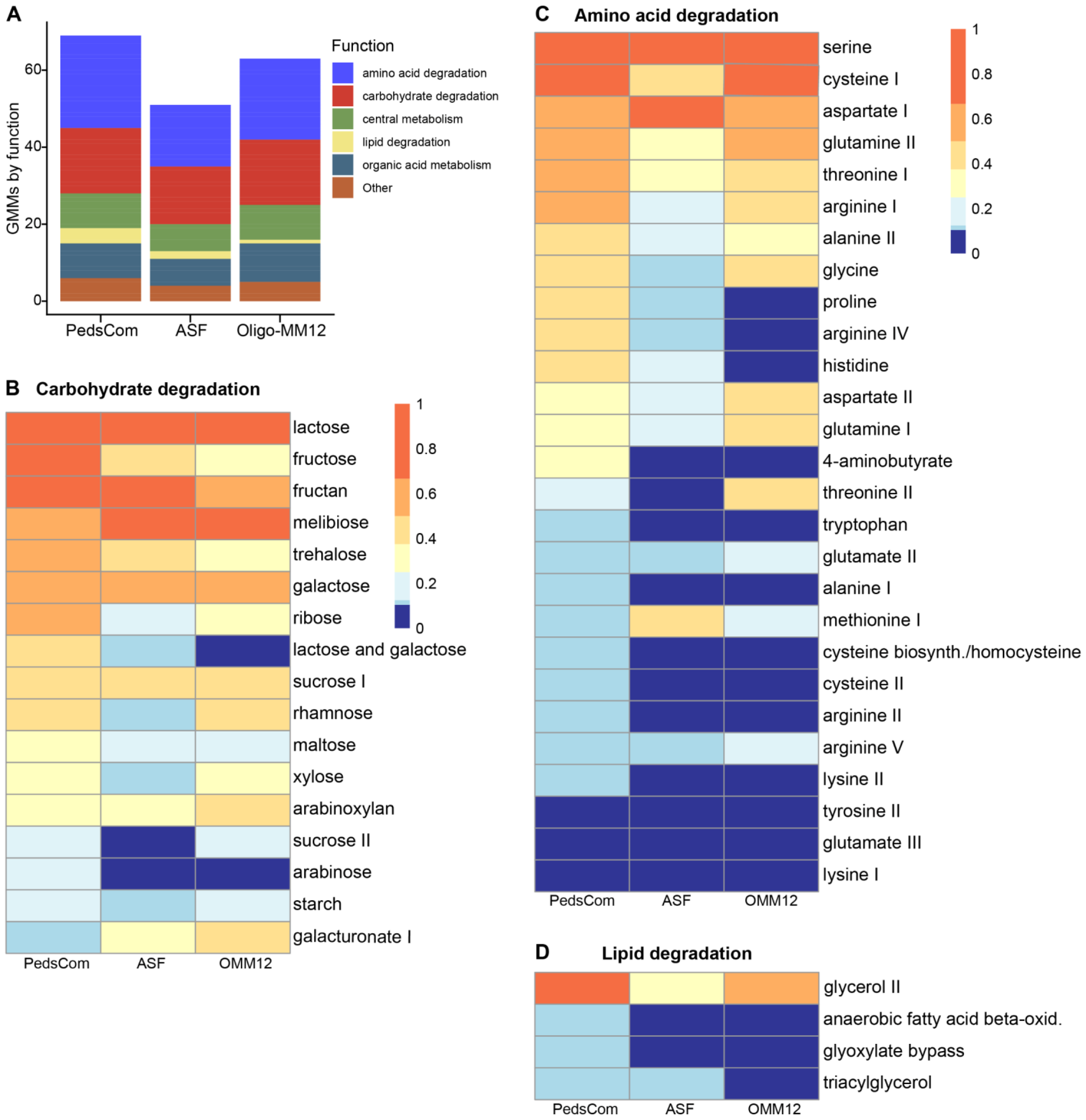
PedsCom consortium contains unique genes associated with pre-weaning microbiomes. **A**. Gut Metabolic Modules (GMMs) found in PedsCom, ASF and Oligo-MM12 isolates. GMMs are considered present if at least 75% of the KEGG orthologs associated with module were found in an isolate. **B**. Heatmap of the proportion of PedsCom, ASF and Oligo-MM12 isolates that contain carbohydrate degradation GMMs. **C**. Heatmap of the proportion of PedsCom, ASF and Oligo-MM12 isolates that contain amino acid degradation GMMs. **D**. Heatmap of the proportion of PedsCom, ASF and Oligo-MM12 isolates that contain lipid degradation GMMs. Heatmap rows are ordered by proportion of PedsCom microbes which contain the module.

For carbohydrates, as expected, PedsCom has a higher proportion of lactose and galactose modules (**Fig. 3B**). However, the carbohydrate degradation modules overall appear similar across the three consortia because the GMM database focuses mostly on mono- and disaccharide degradation capabilities, and not on polysaccharides such as fiber. The PedsCom consortium does have fewer members that can degrade galacturonate and arabinoxylan, constituents of pectin and hemicellulose fibers, respectively.

Amino acid degradation (AAD) GMMs are the major unique functions found in PedsCom (**Fig. 3C**). Of the 27 AAD GMMs present in three consortia, PedsCom has 24. Six AAD GMMs were exclusively found in PedsCom. The most differentially abundant AAD modules in PedsCom relative to ASF and Oligo-MM12 were: γ-aminobutyrate (GABA) (found in 3 of the 9 PedsCom isolates, and 0 of the 8 ASF isolates or 12 Oligo-MM12 isolates), proline (PC = 4/9, ASF = 1/8, OMM12 = 1/12), and histidine (PC = 4/9, ASF = 2/8, OMM12 = 1/12). GABA metabolism and transporters for proline and histidine are significantly associated with the pre-weaning microbiome of human infants^17^. Importantly, the PedsCom members *K. cowanii* (n = 18 GMMs) and *P. distasonis* (n = 15 GMMs) had the most AAD GMMs of any isolates in the three consortia (median per isolate = 6 GMMs).

Lipid degradation modules are also more prevalent within PedsCom (**Fig. 3D**). Of the 13 PedsCom GMMs not found in Oligo-MM12, nine are either amino acid or lipid degradation modules. We observed a similar pattern in PedsCom GMMs absent in ASF, with 10/19 absent functions involved in amino acid or lipid degradation. This pattern indicates that the PedsCom consortium is poised for increased protein and lipid consumption. This skew in PedsCom metabolism is consistent with the functions present in human infant microbiota, highlighting the impact of the milk-based diet on pre-weaning commensal metabolism.

### PedsCom depletes milk-associated amino acids *in vivo*

The higher abundance of amino acid degradation modules in PedsCom relative to Oligo-MM12 led us to hypothesize that there would be lower levels of free amino acids in the gut of PedsCom mice. To test this hypothesis, we performed metabolomic analysis on cecal contents of adult PedsCom- and Oligo-MM12-colonized mice fed standard mouse chow. In support of the hypothesis, we found significantly lower levels of free amino acids in PedsCom mice compared to Oligo-MM12 (24,146 vs 33,278 nmol/g, p<0.05) **(Fig. 4A)**. Hierarchical clustering of amino acid levels in the cecum revealed two clusters of high and low abundance amino acids in the PedsCom gut (**Fig. 4B)**. The top three most abundant free amino acids in milk (glutamic acid, taurine, glutamine)^12^, were significantly depleted in PedsCom cecal contents, relative to Oligo-MM12 colonized mice (**Fig. 4C)**. Additionally, two of the three AAD GMMs (proline, GABA) overrepresented in the PedsCom consortium were significantly reduced in the cecal contents of PedsCom mice. From these analyses, we conclude that PedsCom depletes milk-associated amino acids in the gut, which supports the prediction that PedsCom has a greater capacity to metabolize amino acids.

**Figure 4.**
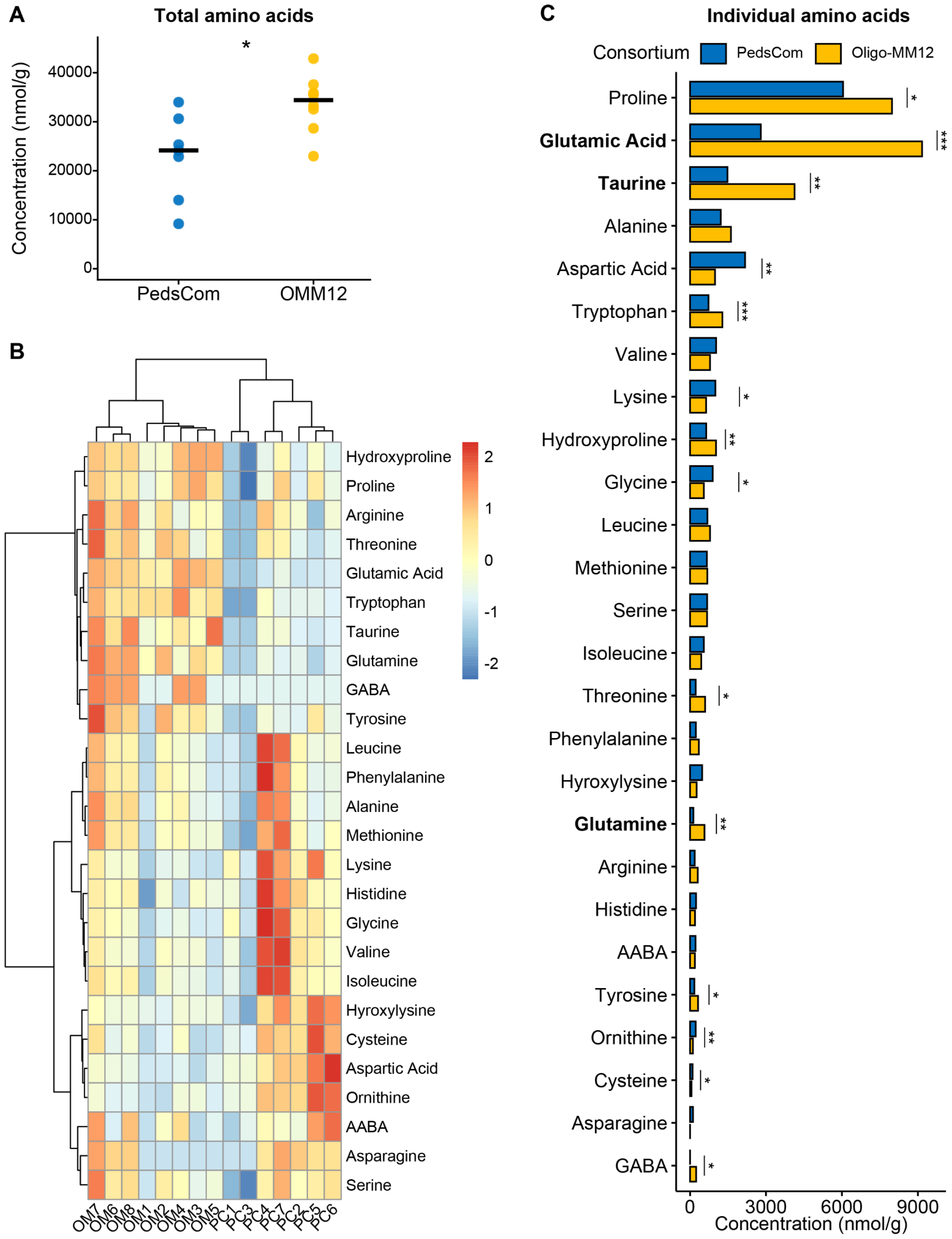
PedsCom colonized mice have reduced amino acid levels in the gut. **A**. Total free amino acid levels (nmol/g) in cecal contents of PedsCom (n = 7) and Oligo-MM12 (n = 8) colonized mice. **B**. Heatmap of free amino acids levels z-score normalized across samples. Hierarchal clustering of columns and rows performed by Ward1 clustering method and Euclidean distance. **C**. Individual amino acid levels (nmol/g) in cecal contents of PedsCom. The 3 most abundant free amino acids found in milk are bolded. Bar plots represent median values. Mann-Whitney-Wilcoxon tests performed, *p<0.05, **p<0.01, ***p<0.001.

## Discussion

We compared the phylogenetic and functional metabolic features of a pediatric gnotobiotic community with two adult communities to better understand the dynamic transition between pre-weaning and post-weaning microbiomes. We made three key observations. First, the PedsCom consortium of microbes is taxonomically and phylogenetically more closely related to organisms found in the infant microbiome than adult-derived gnotobiotic communities. Second, we found a strong predicted metabolic signal for amino acid degradation pathways in the PedsCom community and corroborated this prediction with in vivo metabolomic analysis. Third, the predicted metabolic functions of PedsCom are well-matched to the nutrient profile of milk.

Of these findings, the increase in amino metabolism caught our attention since this was the top difference in predicted metabolic function and amino acid metabolism has the potential to shape the preweaning microbiota and host immune development during the critical weaning transition^34–36^. A sizable proportion (∼25%) of the available nitrogen in milk comes from non-protein sources such as free amino acids^37^. Free amino acids could provide additional nitrogen and energy to pre-weaning gut commensals that is typically unavailable in solid food. This prediction is supported by our data showing that PedsCom colonized adult mice were depleted of free amino acids, particularly those most commonly found in milk, relative to Oligo-MM12 controls. In particular, the presence of *Enterobacteriaceae* member *K. cowanii* provides multiple additional metabolic functions and may be a critical factor in amino acid utilization and metabolism in pre-weaning niche specialization.

Differences in amino acid utilization between gnotobiotic consortia like PedsCom and Oligo-MM12 can have downstream consequences for host cell function and immunity. Several amino acids impacted by PedsCom colonization (glutamic acid, glutamine, GABA) have immunomodulatory and cellular signaling functions in the host gut^38,39^, suggesting that the PedsCom gnotobiotic model can serve as system to study the impact of microbial metabolism on the host during the critical window around weaning.

## Methods

### Mice

Germfree Eα16/NOD mice were housed at the Hill Pavilion gnotobiotic mouse facility, University of Pennsylvania, in flexible film isolators [Class Biologically Clean (CBClean)]. Mice were transferred to sterile cages prior to colonization with the PedsCom and Oligo-MM12 consortia. Mice were fed LabDiet 5021 (Cat# 0006540) and caged on autoclaved beta-chip hardwood bedding (Nepco). Animal studies were approved by the Institutional Animal Care and Use Committee (IACUC) of the University of Pennsylvania.

### Phylogenetic analysis

RNA polymerase subunit beta (RpoB) amino acid sequences of PedsCom, ASF and Oligo-MM12 isolates were aligned with MAFFT (ver. 7.243) using the E-INS-i method^40^. Amino acid substitution models were fitted to the alignment and the L_Gasecuel_2008 with discrete Gamma distribution (LG+G) model with complete deletion was used to generate a maximum likelihood phylogenetic tree of the consortia members in MEGA X^41^.

### Predicted metabolic function

Annotated whole genome sequences of PedsCom isolates were generated as previously described^4^. Genome files for ASF and Oligo-MM12 isolates were obtained from the Bacterial and Viral Bioinformatics Resource Center (BV-BRC). Open reading frames in each isolate were assigned a KEGG Ortholog designation through the KEGG automatic annotation server (KAAS)^42^. Euclidian distance measurements and PCA plots of isolate KOs were generated using the MicrobiomeAnalyst R package^43^. GMM modules^33^ were download thorough github: https://github.com/raeslab/GMMs, and isolate KOs were assigned to a GMM using ANVI’O^44^. A module was considered present if 75% of the genes required for the module were detected in the isolate.

### Intestinal amino acid concentration analysis

PedsCom and Oligo-MM12 isolates were cultured and gavaged into 5–6-week-old germfree C57BL/6J mice as previously described^4^. Colonized mice were housed in sterile cages and sacrificed after two weeks. Cecal contents were collected and flash frozen at -80 °C, prior to metabolomic analysis. Free amino acid concentrations were measured by the PennCHOP microbiome core using the Waters Acuity UPLC system (Cat# 176015000) with an AccQ-Tag Ultra C18 1.7 μm 2.1×100mm column and a photodiode detector array (Waters, Cat# 186003837). Analysis was performed using the UPLC AAA H-Class Application Kit (Waters, Cat# 176002983). Limit of detection for individual amino acids was 1 nmol/g cecal contents.

### Statistical analysis

Heatmaps were generated with the R package pheatmap. Amino acid concentrations were z-score normalized across rows prior to heatmap generation. Statistical analyses were performed in R with the ggpubr package. Significant differences were determined by Mann-Whitney-Wilcoxon tests on median values with p-values < 0.05.

## Acknowledgements

The authors thank Elliot Friedman and Dylan Curry of the PennCHOP microbiome microbial culture and metabolomics core. The authors also thank Joseph Zackular for helpful suggestions on this manuscript. This study was supported by the National Institutes of Health T32 Postdoctoral Program in Clinical Pharmacology T32GM008562 (J.L.), National Institutes of Health grants R01DK133453-01A1 (M.A.S, P.J.P.), and JDRF grant 5-CDA-2020-946-S-B (M.A.S).

## Data availability statement

All data associated with this study is available upon request.

